# Mitochondrial Uncoupling Protein 2 Knockout Promotes Mitophagy to Decrease Retinal Ganglion Cell Death in a Mouse Model of Glaucoma

**DOI:** 10.1101/465153

**Authors:** Daniel T Hass, Colin J Barnstable

**Author notes:** Correspondence: Colin J Barnstable, D. Phil., Department of Neural and Behavioral Sciences, Penn State College of Medicine, Hershey, PA 17033, USA. The authors declare no competing financial interests.

## Abstract

Glaucoma is a neurodegenerative disorder characterized by mitochondrial dysfunction and an increase in oxidative damage, leading to retinal ganglion cell (RGC) death. The oxidative status of RGCs is regulated intrinsically and also extrinsically by retinal glia. The mitochondrial uncoupling protein 2 (UCP2) relieves oxidative and neuronal damage in a variety of neurodegenerative disease models. We hypothesized that deletion of *Ucp2* in either RGCs or retinal glia would increase retinal damage and retinal ganglion cell death in a mouse model of glaucoma. Paradoxically, we found the reverse, and deletion of mitochondrial UCP2 decreased oxidative protein modification and reduced retinal ganglion cell death in male and female mice. This paradox was resolved after finding that *Ucp2* deletion also increased levels of mitophagy in cell culture and retinal tissue. Our data suggest that *Ucp2* deletion facilitates increased mitochondrial function by improving quality control. An increase in mitochondrial function explains the resistance of *Ucp2*-deleted retinas to glaucoma and may provide a therapeutic avenue for other chronic neurodegenerative conditions.

**Significance Statement:** Many unsolved neurodegenerative conditions result from defects in mitochondrial function. Molecular tools that can manipulate mitochondria will therefore be central to developing neuroprotective therapies. Among the most potent regulators of mitochondrial function are the uncoupling proteins, particularly UCP2. In this manuscript we show that while loss of UCP2 does increase mitochondrial membrane potential and the production of reactive oxygen species, it also initiates an increase in mitophagy that is ultimately neuroprotective. This novel protective consequence of uncoupling protein inhibition may lead to new therapeutic approaches to combat neurodegenerative disease, particularly because pharmacological compounds do exist that can selectively inhibit UCP2.

## Introduction

Glaucoma is a group of disorders marked by progressive degeneration and death of retinal ganglion cells (RGCs) that leads to irreversible visual decline. Globally, glaucoma is the most frequent cause of irreversible blindness (Resnikoff and Keys, 2012). Primary open-angle glaucoma (POAG) is the most common form of the disease, and risk for POAG increases with age and elevated intra-ocular pressure (IOP) (Boland and Quigley, 2007; Tham et al., 2014). However, other signs such as a change in retinal morphology or loss of visual function are sufficient for a glaucoma diagnosis without an elevated IOP (Quigley, 2011).

Although glaucoma is often defined by the loss of RGCs, its pathophysiology is multifactorial and involves changes to multiple cell types within the inner retina, including müller glia and astrocytes (Varela and Hernandez, 1997; Carter-Dawson et al., 1998; Kawasaki et al., 2000; Woldemussie et al., 2004). In models of glaucoma, the mitochondria of these cell types become dysfunctional (Lee et al., 2011; Ju et al., 2015). Mitochondrial dysfunction can result in a bioenergetic deficit or increased levels of damaging reactive oxygen species (ROS) in the eye (Tanihara et al., 1997; Varela and Hernandez, 1997; Ferreira et al., 2010). The restoration of mitochondrial homeostasis and anti-oxidative support to both neurons and glia of the retina and optic nerve has become a major focus in research to prevent glaucomatous neurodegeneration (Kong et al., 2009; Lee et al., 2011; Garcia-Medina et al., 2014; Shen et al., 2015; Kimura et al., 2017).

Mitochondrial hydrogen peroxide production is highly dependent on mitochondrial pH and membrane potential (Ψ_m_), and small decreases in Ψ_m_ or pH cause significant decreases in the production of reactive oxygen species (ROS) such as H_2_O_2_ (Korshunov et al., 1997; Miwa et al., 2003). The activity of exogenous agents that decrease Ψ_m_ cannot be easily regulated within cells and their use would risk a complete collapse of Ψ_m_ and thus ATP production. Alternatively, it is the prototypical function of endogenous uncoupling proteins to uncouple the electron transport chain from ATP synthase activity and decrease Ψ_m_ (Fleury et al., 1997; Echtay et al., 2001). The transcription, translation, and activity of endogenous uncoupling proteins are each regulated by multiple factors (Donadelli et al., 2014; Lapp et al., 2014).

There are 5 members of the uncoupling protein gene family (Bouillaud et al., 2001). The mitochondrial uncoupling protein 2 gene (*Ucp2*) is expressed in a variety of tissues, including the central nervous system (Pecqueur et al., 2001; Richard et al., 2001), and codes for a 309 amino acid protein (Fleury et al., 1997). UCP2 activity decreases electron transport chain efficiency, increasing energy expenditure (Zhang et al., 2001) and decreasing ROS generation (Nègre-Salvayre et al., 1997; Arsenijevic et al., 2000). UCP2-mediated uncoupling of mitochondria is cytoprotective under a variety of stressful conditions (Mattiasson et al., 2003; Andrews et al., 2005; Barnstable et al., 2016). As a consequence of *Ucp2* deletion, mice generate more ROS and are increasingly susceptible to cell death following acute exposure to stressors such as the dopaminergic neurotoxin 1-methyl-4-phenyl-1,2,3,6-tetrahydropyridine (MPTP) (Andrews et al., 2005) or focal ischemia (Haines et al., 2010). Notably, Ucp2 deletion can be protective (de Bilbao et al., 2004), deleterious (Andrews et al., 2005; Haines et al., 2010), or have no clear effect on cell survival (Barnstable et al., 2016) in different models of neurodegeneration.

The goal of this study was to determine whether UCP2 normally functions to limit oxidative stress during glaucoma, thereby preventing a more severe form of the disorder. Glaucoma is a disease in which there are greater levels of ROS, and *Ucp2* deletion increases the generation of ROS (Arsenijevic et al., 2000). We therefore hypothesized that decreasing UCP2 levels would increase ROS and RGC death. However, our data suggest that by regulating mitochondrial dynamics, decreases in UCP2 levels can reduce the accumulation of oxidative damage to the retina and decrease retinal ganglion cell death.

## Materials and Methods

### Ethical approval

This study was carried out in accordance with the National Research Council’s Guide for the Care and Use of Laboratory Animals (8th edition). The protocol was approved by the Pennsylvania State University College of Medicine Institutional Animal Care and Use Committee.

### Animals

Wild-type (WT, C57BL/6J) and transgenic mice were housed in a room with an ambient temperature of 25°C, 30-70% humidity, a 12-hr light–dark cycle, and ad libitum access to rodent chow. Transgenic mouse strains, B6;129S-*Ucp2*^*tm2.1Lowl*^/J (*Ucp2*^*fl/fl*^, Stock#: 022394), B6.Cg-Tg(*GFAP-cre/ER*^*T2*^)505Fmv/J (*GFAP-creER*^*T2*^, Stock#: 012849) (Ganat et al., 2006), and Tg(*Thy1-cre/ER*^*T2*^,-EYFP)HGfng/PyngJ *(Thy1-creER*^*T2*^, Stock#: 012708) (Young et al., 2008) were each generated on a C57BL/6J background and purchased from the Jackson Laboratory (Bar Harbor, ME, USA). *Ucp2*^*fl/fl*^ mice contain LoxP sites flanking exons 3 and 4 of the *Ucp2* gene. *GFAP-creER*^*T2*^ and *Thy1-creER*^*T2*^ mice express a fusion product of *cre* recombinase and an estrogen receptor regulatory subunit (*creER*^*T2*^) under the control of the *hGFAP* or *Thy1* promoters. CreER^T2^ activity is regulated by the estrogen receptor modulator and tamoxifen metabolite 4-hydroxytamoxifen (Zhang et al., 1996), and in our studies, cre– mediated recombination of exons 3 and 4 of *Ucp2* was promoted in 1-2 month old mice by daily intra-peritoneal injections of 100 mg/kg tamoxifen (Sigma, T5648) dissolved in sunflower seed oil (Sigma, S5007) for 8 days. *Ucp2*^*fl/fl*^ mice were injected with tamoxifen at the same time points as experimental subjects.

### Breeding Scheme

To produce mice in which *Ucp2* is selectively deleted in *Gfap-* or *Thy1*-expressing cells, *Ucp2*^*fl/fl*^ mice were crossed with *GFAP*-*creER*^*T2*^ or *Thy1-creER*^*T2*^ mice, and *Ucp2*^*fl/+*^; *creER*^*T2*^-positive offspring were crossed with *Ucp2*^*fl/fl*^ mice. The resulting *Ucp2*^*fl/fl*^; *GFAP-creER*^*T2*^ or *Ucp2*^*fl/fl*^; *Thy1-creER*^*T2*^ offspring were bred with *Ucp2*^*fl/fl*^, and the offspring of these pairings were used in this study.

### IOP Measurement

Intra-ocular pressure (IOP) was measured in isoflurane-anesthetized mice using an Icare**®** TonoLab (Icare Finland Oy, Espoo, Finland) rebound tonometer, both before and after injection with polystyrene microbeads. Each reported measurement per mouse per eye is the average of 18 technical replicates. Though the amount of time under anesthesia has been shown to affect IOP (Ding et al., 2011), mice were anesthetized for <10 minutes and yielded stable mean IOP measurements, particularly in control eyes. However, the work of Ding et al., 2011 leads us to believe that our method of measurement underestimates IOP values by at least 3 mmHg, and possibly more in bead-injected eyes (Ding et al., 2011). Mice were only included in this study if their IOP was elevated by 3 mmHg, or if a t-test of their IOP over time between bead and PBS-injected eyes was statistically significant.

### Genotyping

Tissue from ear punches was lysed and digested for genotyping. *Ucp2*^*fl/fl*^ mice were genotyped with PCR primers flanking a LoxP site on the *Ucp2* gene (Table 1, *Ucp2*^*flox*^). To determine whether CreER^T2^-mediated *Ucp2* exon3-4 excision had occurred within a subset of samples, the reverse primer was used together with a primer outside the LoxP-flanked region (Table 1, *Ucp2*^Δ^). PCR conditions to amplify *Ucp2*^*Flox*^ or *Ucp2*^*Δ*^ were as follows: (**1**) 95°C–3 min, (**2**) 95°C–1 min, (**3**) 73°C–1 min, (**4**) 72°C–30 seconds, (**5**) Go to (2) for 15 cycles (−1°C/cycle), (**6**) 95°C–1 min, (**7**) 58°C–1 min, (**8**) 72°C–30 seconds, (**9**) Go to (6) for 20 cycles, (**10**) 95°C–10 min, (**11**) Hold at 4°C. Both the *Thy1-CreER*^*T2*^ and *GFAP-CreER*^*T2*^ genes were genotyped using primers binding to an internal region of *Cre* recombinase (Table 1, Cre). PCR conditions to amplify *Cre* were as follows: (**1**) 95°C–3 min, (**2**) 95°C–1 min, (**3**) 58.1°C–1 min, (**4**) 72°C–30 seconds, (**5**) Go to (2) for 29 cycles, 95°C–10 min, **(6**) Hold at 4°C (Ganat et al., 2006).

**Table 1.**
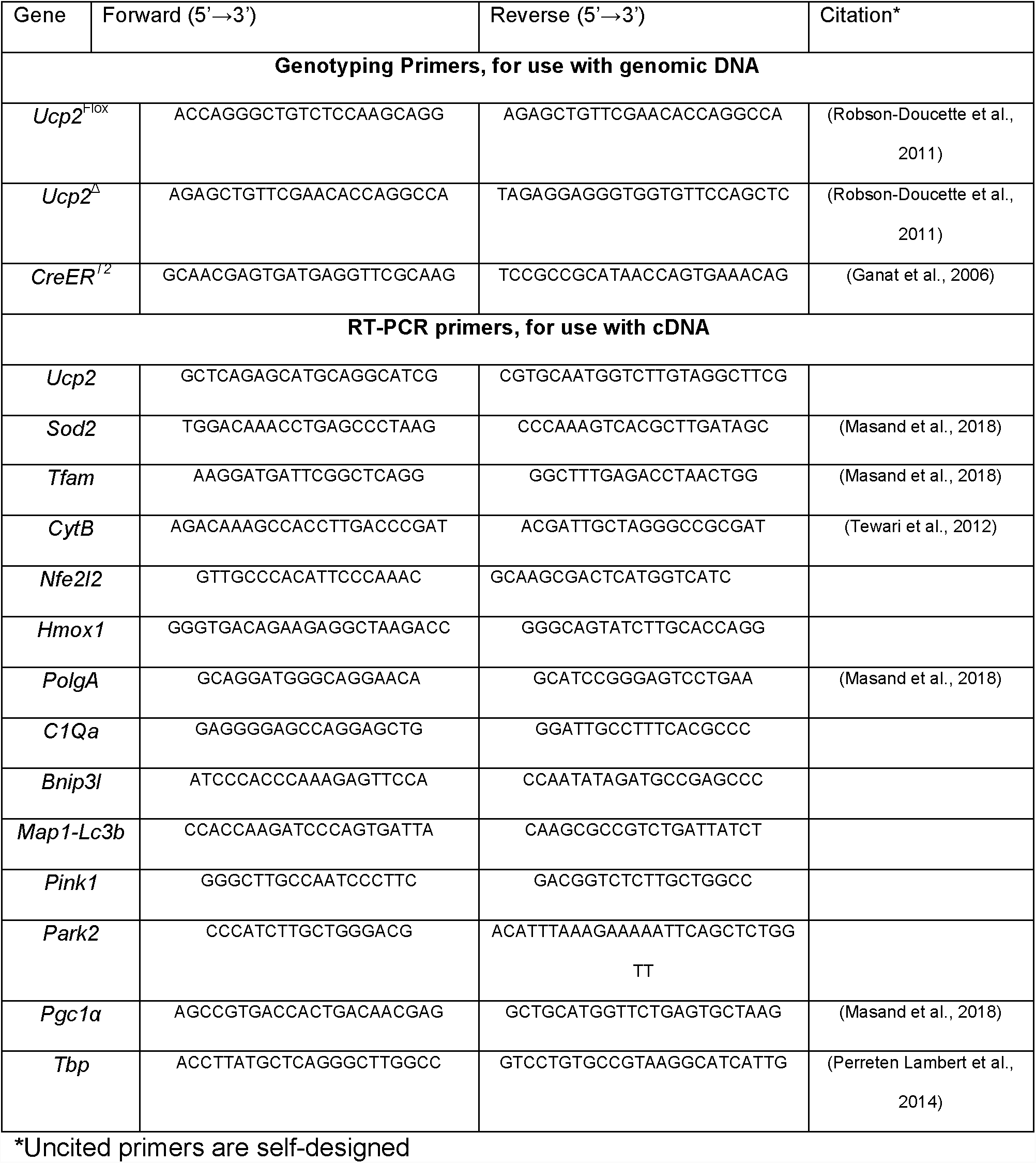
Primers used in this study

### Microbead Injection

We modeled glaucoma in mice by elevating IOP. We increased IOP in 2-4 month old *Ucp2*^*fl/fl*^, *Ucp2*^*fl/fl*^; *GFAP-creER*^*T2*^, and *Ucp2*^*fl/fl*^; *Thy1-creER*^*T2*^ mice of both genders similarly to the Cone et al. ‘4+1’ protocol (Sappington et al., 2010; Cone et al., 2012). At least 24 hours prior to bead injection, we took a baseline IOP measurement to confirm that a given mouse had a ‘normotensive’ IOP. We found that prior to bead injection, IOP is very stable and can be well represented by a single pre-bead measurement (data not shown). We then anesthetized mice with an intraperitoneal (IP) injection of 100 mg/kg Ketamine and 10 mg/kg Xylazine, and treated each eye with topical proparacaine hydrochloride (0.5%) to further anesthetize and hydrate the cornea during injections. We then sterilized 1- and 6-µm diameter polystyrene microbeads (Polysciences, Cat#: 07310-15 and 07312-5) as noted in Cone et al., and estimated bead concentrations on a hemocytometer. We injected 2 µL of 6 µm (at 3×10^6^ beads/µL) and 2 µL of 1 µm (at 1.5×10^7^ beads/µL) microbeads through the cornea using a 50-100 µm cannula formed by a beveled glass micropipette connected by polyethylene tubing to a Hamilton syringe (Hamilton Company Reno, NV, USA). As an internal control, the same volume of sterile 1x phosphate buffered saline (PBS) was injected in to the contralateral eye. We measured post-operative IOP every 3 days for 30 days. Following the terminal IOP measurement, mice were asphyxiated using a Euthanex SmartBox system, which automatically controls CO_2_ dispersion, followed by cervical dislocation. Figure 1B illustrates a visual representation of this experimental design. For measurements of gene expression, protein levels, and mitochondrial function, mice were euthanized 3 days following microbead injection, as IOP is generally maximal or near maximal at this post-operative time point.

**Figure 1.**
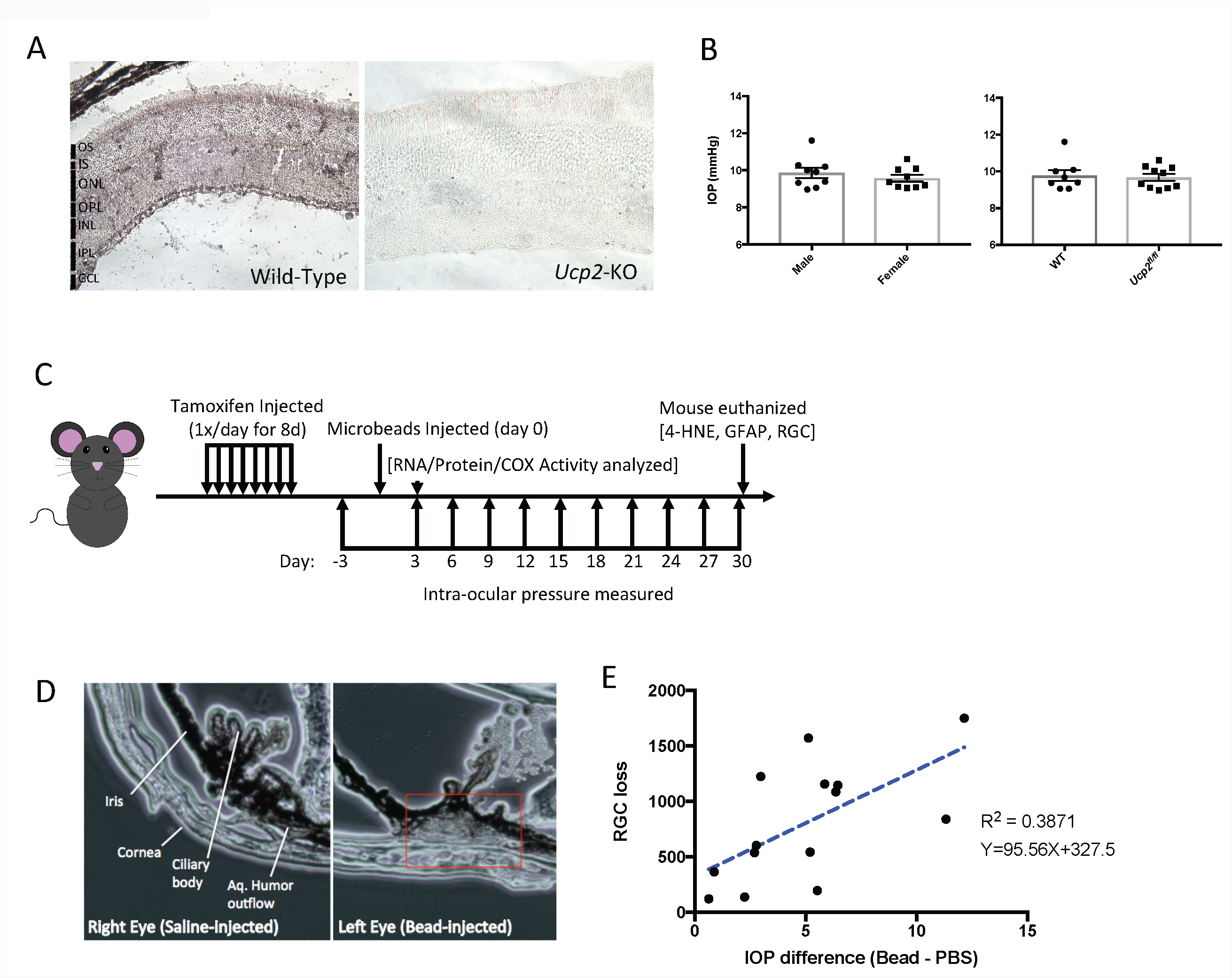
Expression of UCP2 in the retina and characteristics of the microbead model of glaucoma. (**A**) Representative immunohistochemical labeling of UCP2 in retinas from full-body UCP2-KO mice and wild-type mice, showing that UCP2 is distributed throughout the retina, but is expressed to a greater extent in the ganglion cell layer. (**B**) Schematic diagram representing the order and timing of manipulations made to mice prior to and during the microbead model of glaucoma. (**C**) Baseline (pre-bead injection) IOP compared between male and female C57BL6/J mice (left), showing no effect of gender on IOP. These data were pooled and compared against baseline IOP in *Ucp2*^*fl/fl*^ mice. In both cases there is no statistical difference in mean IOP (T-test). (**D**) We verified the location of microbeads following injection through the cornea, and found them packed between the cornea and iris, at the aqueous humor outflow pathway. (**E**) 30 days following IOP elevation, the difference in RGC density between PBS- and bead-injected eyes of Ucp2^fl/fl^ mice was analyzed as a function of average IOP difference between those eyes, showing a positive correlation between IOP elevation and RGC loss.

### Cytochrome-C oxidase Histochemistry

In situ cytochrome-C oxidase (COX) activity was assayed using a modified procedure described by (Wong-Riley, 1979; Murphy et al., 2012). Briefly, following euthanasia, unfixed mouse eyes were harvested, immediately frozen in OCT on dry ice, and cut in to 10 µm sections. Control and experimental samples were frozen and sectioned on the same slide to minimize potential slide-to-slide variability. Slides were incubated in an assay solution containing 4 mM 3’,3’-diamidobenzadine, 100 µM cytochrome C (Sigma, Cat#: 2037), and 5 U/mL catalase (Sigma, Cat#: E3289) for 1h at 37°C. Assay control slides were additionally exposed to 5 mM NaN_3_, which inhibits COX activity. Slides were imaged at 20x magnification. Analysis of staining intensity was performed using the H-DAB setting of the ImageJ color deconvolution tool and subtracting the result from background image intensity.

### Histology and Immunocytochemistry

Immunolabeling of sectioned retinal tissue was performed as previously described (Pinzon-Guzman et al., 2011). Briefly, whole eyes were fixed in 4% paraformaldehyde (Electron Microscopy Sciences, Hatfield, PA, USA) in 1x PBS overnight at 4°C. The next day, eyes were divided in half with a scalpel blade. One half was frozen and sectioned, while the other was labeled as a whole-mount. Frozen tissue were embedded in a 2:1 mixture of 20% sucrose and OCT (Electron Microscopy Sciences), cooled to −20°C, and cut at a 10 µm thickness. Samples for each experiment were located on the same slide to control for assay variability. Prior to immunohistochemical labeling, we unmasked antigens in a pH 6.0 sodium citrate buffer. Sections were permeabilized with 0.2% Triton-X-100 in PBS, blocked with 5% nonfat milk, and incubated in primary antibodies (Table 2) overnight at 4 °C. The following day, sections were washed and incubated in secondary antibody for 3 hours, followed by incubation in 1 µg/mL Hochest-33258 for 20 minutes. After three washes, slides were mounted with 0.5% n-propyl gallate in 1:1 glycerol: PBS. Retinal whole mounts used a similar protocol but with the following modifications: antigens were not unmasked, tissue was incubated in primary antibody for 6 days at 4°C and secondary overnight at 4°C. Labeled tissue was imaged on a Fluoview FV1000 confocal microscope (Olympus). In each experiment, acquisition parameters for a given antibody were held constant. All fluorescent labeling intensity measurements were derived using the ImageJ measurement tool. Unless otherwise specified, intensity measurements were bounded by the ganglion cell layer and the outer nuclear layer. All slides were imaged at 40x magnification, and cells were imaged at 100x magnification. We monitored mitochondrial morphology, size, and number using the MiNa V1 plugin (Valente et al., 2017). We measured colocalization between LC3B and TOMM20 using the ImageJ plugin “Coloc 2” and reporting the Manders’ overlap coefficient corresponding to the TOMM20 image channel. Tissue LC3B puncta were detected using the “Find Maxima…” tool with a noise tolerance of 1000, which of all the methods tested best recapitulates a visual search for LC3B puncta.

**Table 2.**
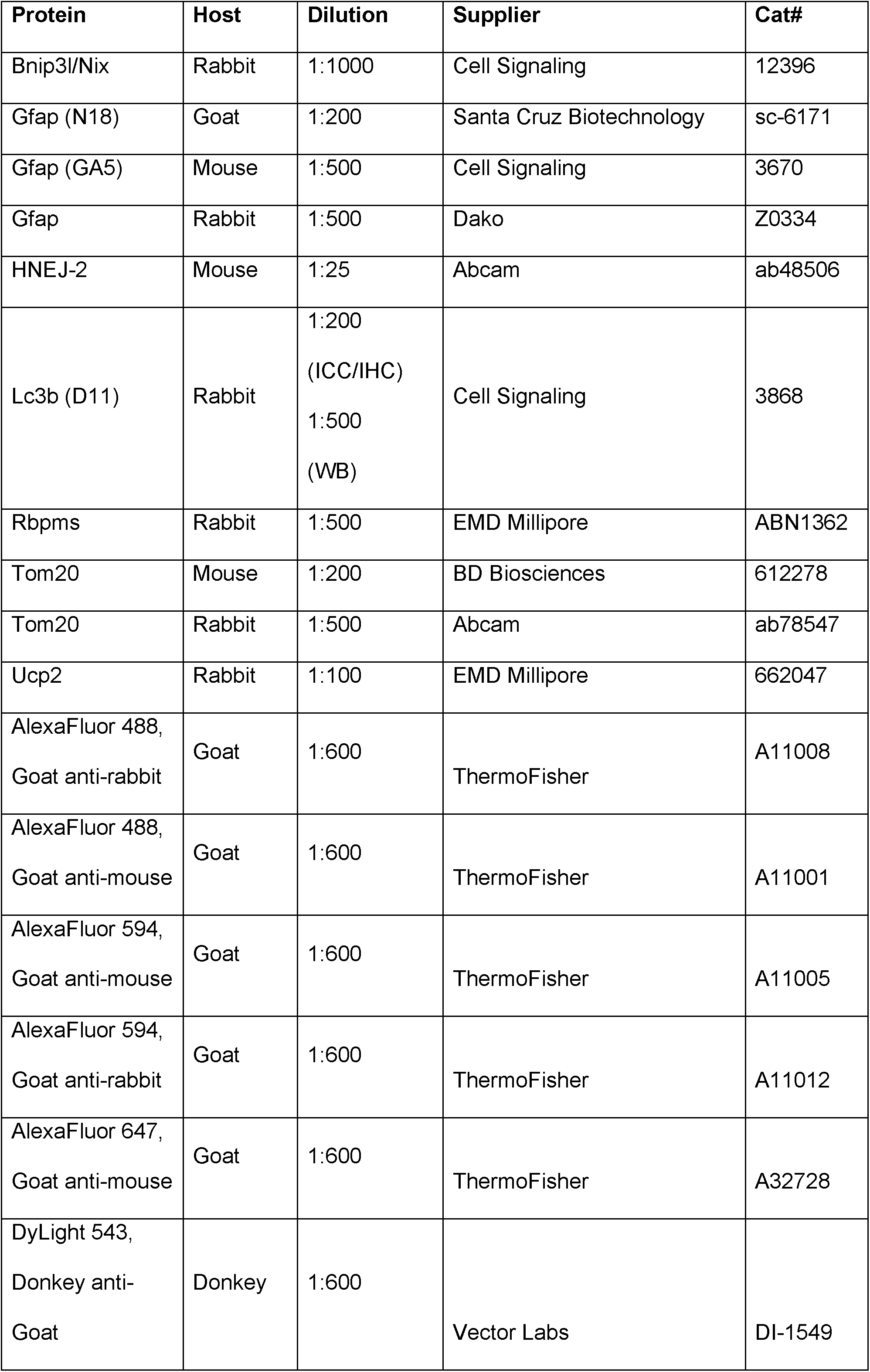
Antibodies used in this study

### Retinal Ganglion Cell Quantification

RGC density was estimated in the ganglion cell layer of RBPMS immuno-labeled retinal whole-mounts (Rodriguez et al., 2014). We used RBPMS due to the high concordance between RGC soma labeling with this marker and with several other well-characterized markers, both normally (Rodriguez et al., 2014) and pathologically after optic nerve damage (Kwong et al., 2011). This concordance suggests that loss of RBPMS labeling was due to loss of RGCs and not to loss of RBPMS expression. For each retina, RBPMS+ cells were counted across 3-4 well-separated fields, with each field measuring 317.95 µm x 317.95 µm and centered 1000 µm from the optic nerve head. This corresponds to a total area of 0.3-0.4 mm^2^ counted per retina. Cell counts were converted to measurements of RGC density and averaged for a single retina. The observer counting RGC numbers in these images was blinded to the identity of each sample. RGC density was calculated from cell count these images, and RGC loss was calculated as the difference in these averages between PBS injected retinas and contralateral Bead-injected retinas. Mean±SEM and median RGC densities were 4283±83 and 4169 cells/mm^2^, respectively in PBS-injected *Ucp2*^fl/fl^ retinas, comparable to values found in the literature for this marker (Rodriguez et al., 2014; Wang et al., 2017). The percentage of RGC death following microbead injection was also similar to our own findings using a different RGC marker, Brn3a (Hass et al., unpublished observations). We did not find a significant effect of bead-injection on retinal quadrant RGC density, and our images were therefore averaged from across all retinal quadrants for a given sample.

### RNA Isolation and Quantitative Real-Time PCR

Frozen tissue was lysed in TRIzol (Thermo-Fischer, Cat#: 15596018) and RNA precipitated using the manufacturer’s recommended procedure. Final RNA concentration was measured using a NanoDrop ND-1000 Spectrophotometer prior to reverse transcription. We reverse transcribed RNA using SuperScript III (Thermo-Fischer, Cat#: 18080093) with random hexamers. cDNA was amplified with iQ SYBR Green Supermix (Bio-Rad, Cat#: 1708882) and quantitated on a Bio-Rad iCycler. All primers designed to measure gene expression (Table 1) do not amplify genomic sequences, and qPCR reactions were followed by a melt curve analysis to confirm the amplification of a single PCR product. All results were normalized against the TATA-box binding protein (*Tbp*) cDNA using the ΔΔC_t_ method. *Tbp* is stably expressed during both mouse retinal development (Adachi et al., 2015) and in retinal endothelial cells treated with innate immune system stimulants (Wei et al., 2013).

### Primary Astrocyte Culture

Primary mouse cortical astrocytes were isolated from postnatal day 1-4 mice as previously described (Sarafian et al., 2010; Lapp et al., 2014). Briefly, mice were decapitated and brains were removed from the skull. In tissue culture medium, a ∼1 cm portion of superior cerebral cortex was pinched off of the brain using curved forceps. Meninges were removed, and the tissue was triturated with a sterile flame-polished glass Pasteur pipette until it formed a single cell suspension, approximately 20x. The suspension was filtered through a 70 µm cell strainer to remove larger debris, centrifuged at 500 x g and 4°C for 5 min, resuspended in growth medium (Dulbecco’s Modified Eagle’s Medium/Ham’s F12 supplemented with 2 mM L-glutamine, 15 mM HEPES, 10% fetal bovine serum, and 10 ng/mL gentamicin), and plated in a T-25 tissue culture flask. Cells were grown at 37°C in a 5% CO_2_/balance air atmosphere. After the cells have reached confluence between 7-14 days in vitro (DIV), contaminating cells were shaken off at 250 RPM overnight and growth medium was replaced. Astrocyte-enriched cultures were then passaged at least 48 hours following medium replacement for experimentation, and used for experimentation at passage #2 or 3. All cells used in this study were exposed to 1 µM 4-hydroxytamoxifen (Sigma, Cat#: H6278) for 24 hours prior to studies of *Ucp2* function.

### Measurement of Mitochondrial Membrane Potential and Oxidative Status

We determined mitochondrial membrane potential (Ψ_m_) and relative oxidative status were determined on a fluorescence-enabled microplate reader (BioTek Synergy II) using black tissue-culture-treated 96-well plates (Corning, Cat#3603) seeded with 30,000 cells/well. We treated all primary astrocytes from *Ucp2*^*fl/fl*^ and *Ucp2*^*fl/fl*^; *GFAP-creER*^*T2*^ mice with 4-hydroxytamoxifen and washed it out at least 24 hours prior to the assay. To measure membrane potential, we loaded cells with 5,5’,6,6’-tetrachloro-1,1’,3,3’-tetraethyl-benzimidazolylcarbocyanine iodide (JC-1; 2 µg/mL; Thermo-Fischer, Cat#: T3168) in culture medium for 30 min. We measured red and green fluorescence 5 minutes after washing out each probe. Each assay contained controls treated with the membrane permeant protonophore Carbonyl cyanide-*4*-(trifluoromethoxy) phenyl-hydrazone (FCCP, 1 µM, Caymen Chemical). We determined the effect of UCP2 on oxidant production using the redox-sensitive probe Chloromethyl–2′,7′-di-chlorofluorescein diacetate (CM-H_2_-DCFDA, 40 µM; Thermo-Fischer, Cat#: C6827) in 1x PBS supplemented with 1 mM glucose and 2 mM GlutaMax (Thermo-Fischer, Cat#: 35050-061). The probe was washed out following a 30-minute incubation. We measured oxidant production in the plate reader’s kinetic mode, which took serial measurements of CM-H_2_-DCFDA fluorescence over time. The increase in fluorescence (ΔF) over 30 minutes was divided by initial fluorescent intensity (F_0_). This rate of increase in CM-H_2_-DCFDA fluorescence was normalized to the mean ΔF/F_0_ of *Ucp2*^*fl/fl*^ cells to minimize inter-assay variability.

### Extracellular Flux Analysis

Oxygen consumption (OCR) and extracellular acidification rate (ECAR) were used as proxies of oxidative and glycolytic metabolism, respectively. We performed a ‘mitochondrial stress test’ on cultured astrocytes (seeded at 30,000 cells/well) using inhibitors of electron transport at the indicated final concentrations: oligomycin (25 µg/mL; Caymen Chemical), FCCP (1 µM), antimycin A (1 µM), and rotenone (1 µM). Following the assay, we measured DNA concentration in each well for normalization of OCR traces.

### Western blot analysis

Western blot analysis was preformed as previously described (Pinzon-Guzman et al., 2011). Briefly, Dissected retinal tissue was homogenized in RIPA buffer supplemented with Halt protease inhibitor cocktail (Thermo-Fischer, Cat#: 78430) and a phosphatase inhibitor cocktail set II (EMD-Millipore, Cat#: 524625). Protein concentration was estimated with a DC protein assay (BioRad, Cat#: 5000111). 20 µg retinal homogenate was loaded on a 4-12% Criterion TGX polyacrylamide gel (Bio-Rad, Cat#: 5678124) and run at 240 V for 35 minutes prior to semi-dry transfer to a 0.45 µm thick nitrocellulose membrane (Bio-Rad, Cat#: 1620115) at 16 V for 40 minutes. Leftover protein was visualized with Coomassie Blue staining, and the sum of Coomassie–labeled histone bands were used for protein normalization.

The membrane was washed in 1x Tris buffered saline with 0.1% Tween-20 (TBS-T) and blocked in TBS-T with 2% (w/v) blotting grade blocker (Bio-Rad, Cat#: 1706404). Membranes were incubated in primary antibodies (Table 2) diluted in blocking buffer overnight at 4°C. Membranes were washed in TBS-T and incubated in secondary antibody for 2 hours at room temperature prior to generation of chemiluminescent bands with Pierce ECL Western Blotting Substrate (Thermo-Fischer, Cat#: 32106) and visualization on photosensitive films (Thermo-Fischer, Cat#: 34091). Band density was determined from scanned developed films using ImageJ.

### Statistical analysis

We performed all statistical analyses in GraphPad Prism. Linear trends were analyzed with a linear regression. We determined the statistical effect of one independent variable on 2 groups used a Student’s t-test or paired sample t-test in cases where samples were matched (e.g. the control was the contralateral eye of the same animal). We analyzed the effect of one variable on >2 groups using a one-way ANOVA with a Holm-Sidak post-hoc analysis. We analyzed the effect of 2 variables using a 2-way ANOVA with a Holm-Sidak post-hoc analysis. The statistical significance threshold was p<0.05 for all tests.

## Results

### UCP2 Localization in the Retina and Characteristics of the Microbead Model

To confirm that UCP2 is normally present in the retina and to determine its cellular localization, we immunolabeled fixed retinal tissue from wild-type C57BL6/J and whole-body *Ucp2* knockout mice (courtesy of Dr. Sabrina Diano). Retinas from *Ucp2*-knockout mice were used as a control for antibody specificity. UCP2 was distributed throughout the mouse retina, with strong expression within the retinal ganglion cell layer (GCL) and photoreceptor inner segments (left panel, Fig. 1A). In order to study the actions of UCP2 in a mouse model of glaucoma, we implemented a microbead model that elevates intra-ocular pressure (IOP), illustrated in Figure 1B. Prior to further studies, we confirmed that neither gender nor genotype had an effect on the baseline IOP characteristics of the mouse eye (Fig. 1B). We also confirmed that injection of microbeads in to the anterior chamber of the eye resulted in their migration to the aqueous humor outflow pathway, at the irido-corneal angle (Fig. 1D). We found a significant (p=0.0175, df=12, n=14) but overall weak relationship (R^2^=0.39) between the bead-induced increase in IOP and a decrease in RGC survival, measured by a decrease in the density of RBPMS+ retinal ganglion cells in retinal whole-mounts between PBS- and microbead-injected eyes (Fig. 1E).

### Elevated IOP Decreases Mitochondrial Function And Increases Oxidative Stress

One stable mark of oxidative damage is accumulation of lipid peroxidation products such as 4-hydroxy-2-nonenal (HNE) (Malone and Hernandez, 2007). HNE can modify histidine residues on proteins, forming a stable adduct that we used as a proxy for oxidative stress in tissue. To determine whether retinas are stressed in the microbead glaucoma model, we compared HNE labeling in the retinal layers of microbead- and PBS-injected eyes. Microbead-injection significantly increased HNE reactivity in the inner retina to 159±29% of contralateral PBS-injected eyes (p=0.0035, df=36, n=5), but not in the outer retina (98±23% of contralateral control). Analysis of individual inner retinal layers showed increases in HNE but none of these reach significance following adjustment for multiple comparisons, suggesting that although the effects of IOP on oxidative stress may occur in multiple cell types, its main effect is within the cell type that degenerates in glaucoma (Fig 2A-B). To determine the effect of IOP elevation on mitochondrial function, we tested cytochrome oxidase (COX) activity in fresh-frozen slices of retina from PBS- and Bead-injected mice and found that by 3 days post bead-injection, there was a significant decrease in retinal cytochrome oxidase activity (p=0.0313, df=4, n=5, Fig. 2C-D). The decrease in COX activity was associated with expression of oxidative stress and glaucoma-related genes. We measured *Hmox1, Nfe2l2, Ucp2, Sod2*, and *C1Qa* in retinas from a group of untreated (naïve) mice, and from PBS-injected, and microbead-injected mice, 3 days following bead injection (n=3, df=6). Bead-injection increased the expression of *Hmox1* (856±294%, p=0.0217), *Nfe2l2* (149±14%, p=0.033), and *C1Qa* (728±295%, p=0.048) relative to contralateral bead-injected retinas, and caused smaller, non-significant increases in *Sod2* (117±35%, p=0.73) and notably in *Ucp2* as well (366±196%, p=0.29, Fig. 2E). In all cases where we measured gene expression, there was no notable or statistically significant difference between retinas of naïve and PBS-injected mice of the same genotype. In all subsequent experiments we used contralateral PBS-injected retinas as a control for microbead-induced IOP elevation, and consider it useful as a proxy for normal retinas.

**Figure 2.**
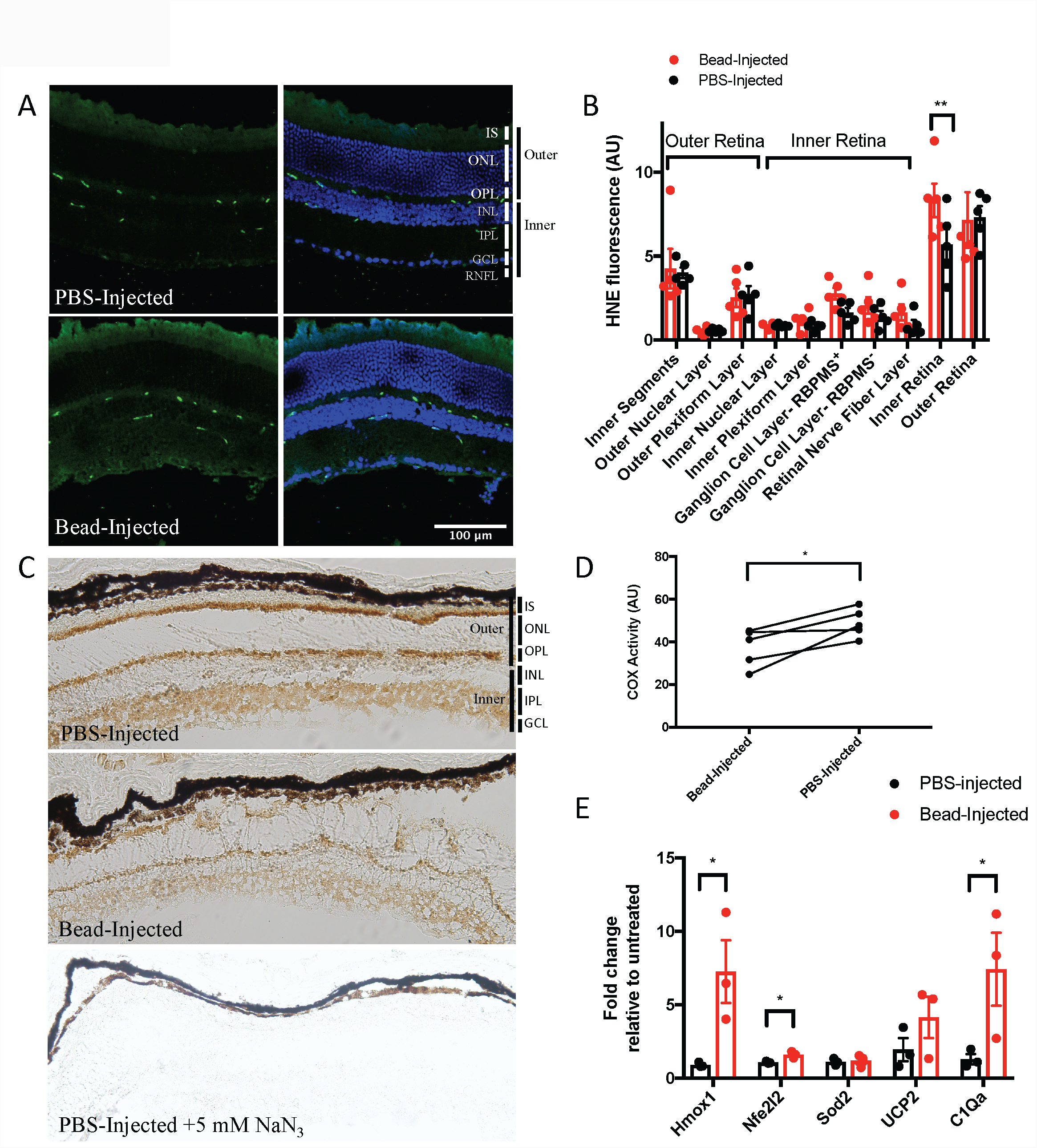
Elevating intra-ocular pressure increases oxidative stress and decreases mitochondrial function. **(A)** 4-hydroxynonenal (green) and Hoechst 33258 (blue) labeling of retinal sections from PBS- and microbead-injected eyes, 30 days post-injection. Scale bar = 100 µm. (**B**) Quantification of immunoreactivity in different retinal layers reveals that HNE labeling is significantly greater within the inner retina of bead injected eyes, relative to contralateral PBS-injected control eyes (black bars=PBS-injected, red bars=bead-injected, n=5). (**C**) COX histochemistry of retinal sections from PBS- and microbead-injected eyes 3 days post-injection, showing that bead injection appears to decreases cytochrome C oxidase activity (DAB labeling) early in glaucoma (n=5). (**D**) Quantification of retinal DAB labeling from (C), related to cytochrome oxidase activity (**E**) Gene expression changes early in glaucoma show that (I) PBS-injection does not appear to significantly alter gene expression relative to untreated retinas, and (II) bead-injection increases the expression of several antioxidant and glaucoma-related genes, including Hmox1, Nfe2l2, and C1Qa (n=3). IS=Inner segments, ONL=outer nuclear layer, OPL=outer plexiform layer, INL=inner nuclear layer, IPL=inner plexiform layer, GCL=ganglion cell layer, RNFL=retinal nerve fiber layer, Outer=outer retina, Inner=inner retina. *=p<0.05, **=p<0.01.

### *Ucp2* Deletion Increases Mitochondrial Membrane Potential (Ψ_m_) and Oxidative Stress

To investigate the effects of *Ucp2* deletion on cellular function, we used primary cortical astrocytes cells from *Ucp2*^*fl/fl*^ and *Ucp2*^*fl/fl*^; *GFAP-creER*^*T2*^ mice. In figure 3A (II) we show a schematic diagram of the floxed *Ucp2*, including the region excised by cre recombinase, and the PCR primers used to determine genotype or cre recombination. We also show the two different variants of cre used in this study (I) as well as (III) a schematic indicating which figures use primary cells and which used retinal tissue. In figure 3B we show a representative agarose gel from 4-hydroxytamoxifen-pretreated *Ucp2*^*fl/fl*^ (Fig. 3B, lanes 1-2) and *Ucp2*^*fl/fl*^; *GFAP-creER*^*T2*^ astrocytes (lanes 3-4), from which we amplified a PCR product (Ucp2^Δ^) that is only detected in cre-expressing cells, following cre-mediated *Ucp2* exon3-4 excision. We determined the effect of *Ucp2* on mitochondrial transmembrane potential (Ψ_m_), oxidative stress, and cell respiration using these astrocytes. We measured Ψ_m_ with the vital dye JC-1, which equilibrates in mitochondria as red fluorescent aggregates and outside as green fluorescent monomers. The ratio of aggregate/monomers is proportional to Ψ_m_, and we found that *Ucp2* deletion (n=7) increased Ψ_m_ to 120±7% of Ucp2^fl/fl^ (n=7) cells (p=0.013, df=15). As a control, we applied the membrane-permeant protonophore FCCP (1 µM, n=3), which reduced Ψ_m_ to 72±4% of controls (p=0.013, df=15, Fig. 3C). Oxidative stress was measured using CM-H_2_-DCFDA, a cell permeant probe that becomes fluorescent upon oxidation, and we measured the rate (ΔF/F_0_) of probe oxidation over 30 minutes. Relative to *Ucp2*^*fl/fl*^ controls, *Ucp2* deletion increased oxidative stress to 133±12% (p=0.0452, df=16, n=7). As a control to verify assay function, *Ucp2*^*fl/fl*^ cells were treated with 4 µM Antimycin A, which non-significantly increased oxidative stress to 122±10% of controls (p=0.13, df=16, n=5, Fig 3D). We measured oxygen consumption rate using a seahorse extracellular flux assay in *Ucp2*^*fl/fl*^ and *Ucp2*^*fl/fl*^; *GFAP-creER*^*T2*^ astrocytes. Following each assay we normalized respiratory activity to DNA concentration. Basal respiration (untreated) over time was greater in *Ucp2*^*fl/fl*^; *GFAP-creER*^*T2*^ cells (p<0.0001, df=1, n=3), as was maximal respiration (p=0.0004, df=1, n=3), suggesting unexpectedly greater mitochondrial function in cells lacking a functional copy of *Ucp2* (Fig. 3E).

**Figure 3.**
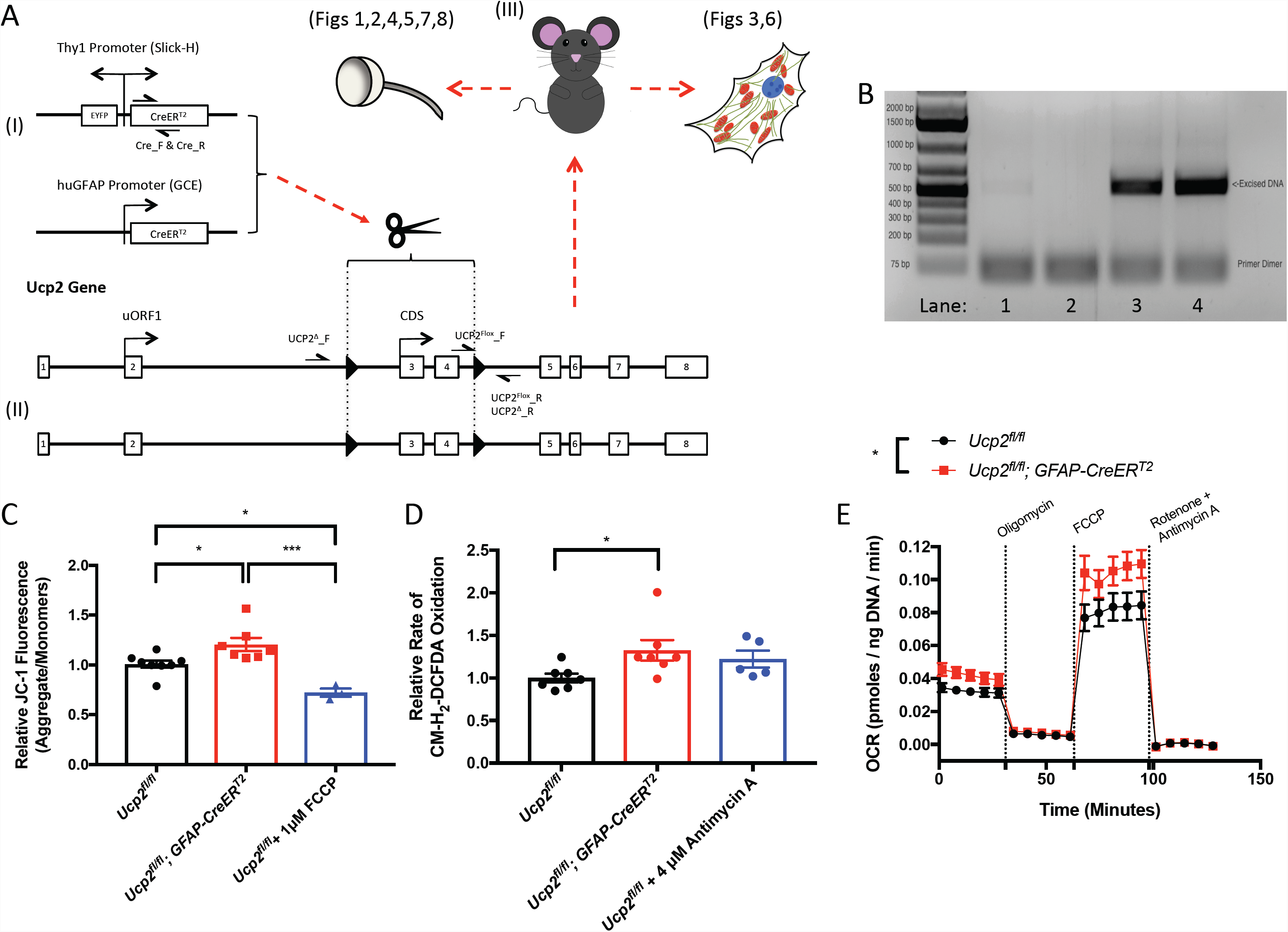
*Cre*-mediated *Ucp2* excision increases Ψ_m_, the rate of oxidant generation, and mitochondrial oxygen consumption in primary astrocytes. (**A**) Schematic diagram of transgenic mouse genotypes for mice used in this study, the approximate primer locations for genotyping these animals, the region of DNA excised by Cre recombinase activity, tissues used from these mice, and the figures in which these tissues are used. (**B**) Example genotyping PCR of 4-hydroxytamoxifen pretreated astrocytes from *Ucp2*^*fl/fl*^ (lanes 1-2) and *Ucp2*^*fl/fl*^; *GFAP-creER*^*T2*^ (lanes 3-4) mice. The primers in this reaction only amplify when the >1000bp LoxP-flanked region of the Ucp2 gene is removed. (**C**) The JC-1 red/green fluorescence ratio in these primary astrocytes (n=8) in increased by UCP2 deletion (n=7) and decreased by 1µM FCCP (n=3), revealing that UCP2 normally functions to decrease Ψ_m_ in these cells. (**D**) The rate of CM-H_2_-DCFDA oxidation is also increased with *Ucp2* deletion (n=7) or with 4µM Antimycin A (n=5). (**E**) Seahorse XF96 measurement of cellular oxygen consumption demonstrating that *Ucp2* deletion increases the basal and maximal mitochondrial oxygen consumption of astrocytes, though not uncoupled respiration (n=3). *=p<0.05, ***=p<0.005.

### *Ucp2* Deletion Decreases Retinal Ganglion Cell Loss During Glaucoma

Cre-mediated *Ucp2* deletion was confirmed by measuring retinal transcript levels, which decreased to 54±14% and 58±13% in *Ucp2*^*fl/fl*^; *GFAP-creER*^*T2*^ and *Ucp2*^*fl/fl*^; *Thy1-creER*^*T2*^ mice relative to *Ucp2*^*fl/fl*^ controls (Fig. 4A). These extents of Ucp2 decrease were close to the expected value, as Ucp2 transcript appears to be greatly expressed within cells of the inner retina (Fig. 1A). Using reporter mice in which EGFP or YFP are co-expressed with *cre* recombinase, we confirmed the localization of these *cre* variants to retinal glia or ganglion cells (Fig. 4B). We injected beads in to mice with *Ucp2* selectively deleted in *Gfap-* or *Thy1*-expressing cells. On average, bead injection increased mean IOP by 9.50±1.49 mmHg in *Ucp2*^*fl/fl*^ mice, 3.78±0.63 mmHg in *Ucp2*^*fl/fl*^; *GFAP-creER*^*T2*^ mice, and 8.28±1.71 mmHg in *Ucp2*^*fl/fl*^; *Thy1-creER*^*T2*^ mice (Fig. 4C). In addition to our own data (Fig. 1E) others have demonstrated that the extent of IOP increase in the microbead model does not always correlate well with the extent of RGC loss (Cone et al., 2010), so differences in the extent of IOP increase between *Ucp2*^*fl/fl*^, *Ucp2*^*fl/fl*^; *Thy1-creER*^*T2*^, and *Ucp2*^*fl/fl*^; *GFAP-creER*^*T2*^ mice should not significantly alter the extent of retinal ganglion cell death. To determine the effect of *Ucp2* deficiency on glaucomatous RGC death, we counted the density of RBPMS+ RGCs on whole-mount retinas from *Ucp2*^*fl/fl*^, *Ucp2*^*fl/fl*^; *GFAP-creER*^*T2*^ and *Ucp2*^*fl/fl*^; *Thy1-creER*^*T2*^ retinas (Fig. 4D). We determined the effect of glaucoma on RGC density using a 2-way ANOVA with Genotype and Bead-Injection as independent variables. A post-hoc analysis revealed significant differences in bead-induced RGC loss between *Ucp2*^*fl/fl*^ retinas (607±148 cells/mm^2^ in, df=49, n=29), *Ucp2*^*fl/fl*^; *GFAP-creER*^*T2*^ (−201±196 cells/mm^2^, p=0.0039, n=10), and *Ucp2*^*fl/fl*^; *Thy1-creER*^*T2*^ retinas (−169±151 cells/mm^2^, p=0.0039, n=13, Fig. 4D-E). These data suggest that surprisingly, reduced retinal UCP2 levels decreased RGC loss following IOP elevation.

**Figure 4.**
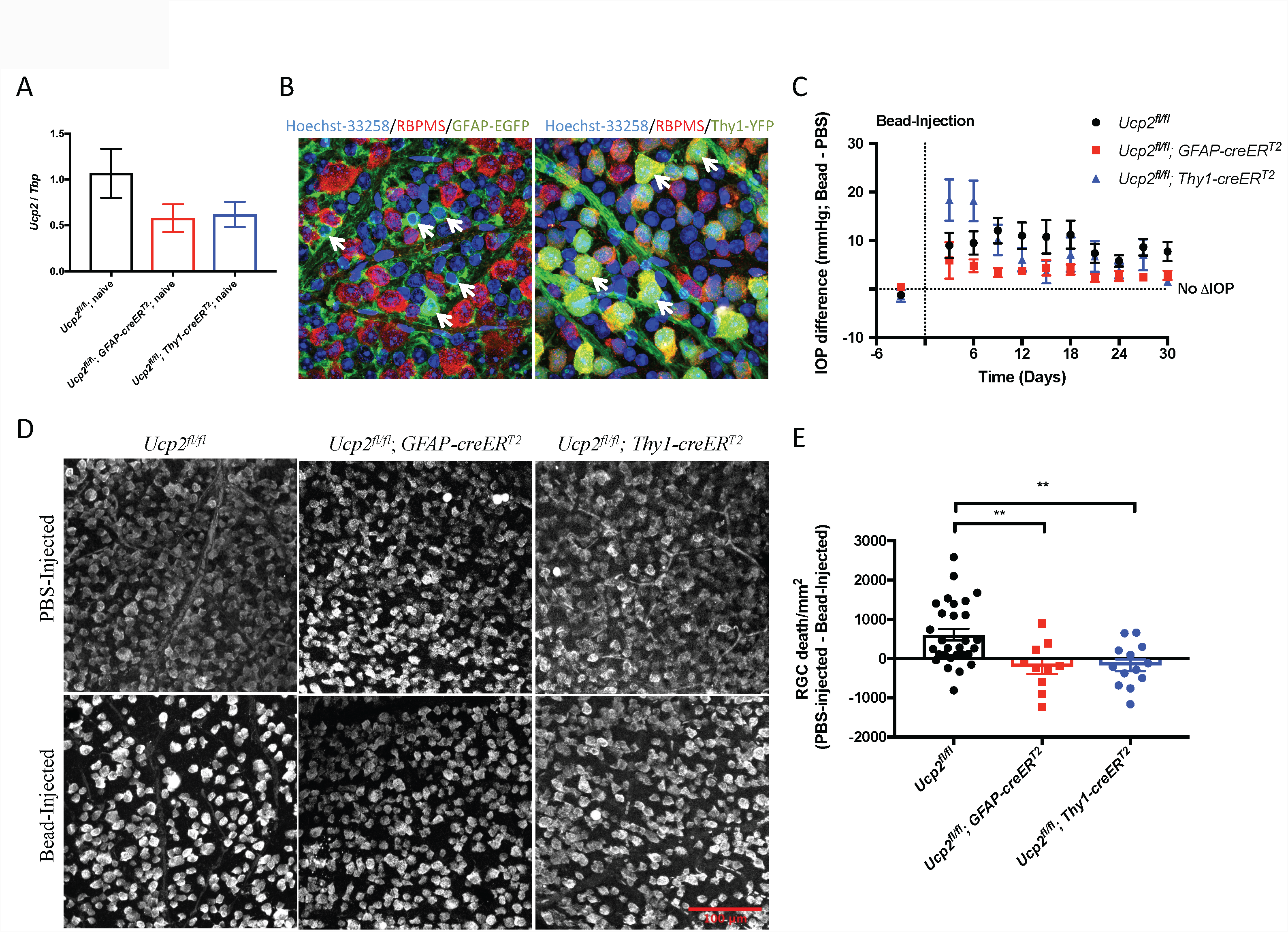
*Ucp2* deletion in either glia or RGCs decreases RGC loss in glaucoma. (**A**) *Ucp2* expression in U*cp2*^*fl/fl*^, *Ucp2*^*fl/fl*^; *GFAP-creER*^*T2*^, and *Ucp2*^*fl/fl*^; *Thy1-creER*^*T2*^ retinas, showing a decrease in *Ucp2* expression in these transgenic mice (**B**) Whole-mount retinas labeled with the nuclear label Hoechst-33258 (blue), the retinal ganglion cell marker RBPMS (red), and the endogenous fluorescence of EGFP in *GFAP-creER*^*T2*^ mice (left) or YFP in *Thy1-creER*^*T2*^ mice (right). (**C**) IOP difference between microbead and PBS-injected *Ucp2*^*fl/fl*^, *Ucp2*^*fl/fl*^; *GFAP-creER*^*T2*^, and *Ucp2*^*fl/fl*^; *Thy1-creER*^*T2*^ eyes over time following a single microbead injection. (**D**) Example RBPMS-labeled RGCs in PBS- and Bead-injected eyes from *Ucp2*^*fl/fl*^ (n=29), *Ucp2*^*fl/fl*^; *GFAP-creER*^*T2*^ (n=10), and *Ucp2*^*fl/fl*^; *Thy1-creER*^*T2*^ (n=13). Scale bar (red) = 100 µm. Differences in RGC density between PBS- and Bead-injected eyes are quantified in (**E**). A difference of “0” indicated no retinal ganglion cell death. **=p<0.01.

### The Mechanism Of Ucp2-Deletion Mediated Neuroprotection

To determine the effect of UCP2 on retinal oxidative stress, we measured HNE immunoreactivity in retinas of bead-injected eyes from *Ucp2*^*fl/fl*^, *Ucp2*^*fl/fl*^; *GFAP-creER*^*T2*^, and *Ucp2*^*fl/fl*^; *Thy1-creER*^*T2*^ mice (Fig. 5A-B). *Ucp2*^*fl/fl*^; *Thy1-creER*^*T2*^ (n=8) and *Ucp2*^*fl/fl*^; *GFAP-creER*^*T2*^ (n=7) accumulated significantly less HNE (p=0.02 and 0.03 respectively, df=20) than *Ucp2*^*fl/fl*^ controls (n=8), providing a quantitative proxy for the qualitative finding that the inner retina was less oxidatively damaged (Fig. 5B). Increases in retinal GFAP are a sign of Müller cell reactivity, and in *Ucp2*^*fl/fl*^ mice, bead injection increased GFAP intensity to 239% of control (p=0.0085, df=61, n=13-23). In *Ucp2*^*fl/fl*^; *GFAP-creER*^*T2*^ or *Ucp2*^*fl/fl*^; *Thy1-creER*^*T2*^, bead injection did not significantly increase our semi-quantitative measure of GFAP labeling intensity, though there was a trend toward increase for each genotype (Fig. 5C-D). This is visualized in representative retinal sections of bead-injected transgenic mice, within which Gfap+ müller cell fibers are detectable but reduced in overall intensity relative to controls (Fig 5C).

**Figure 5.**
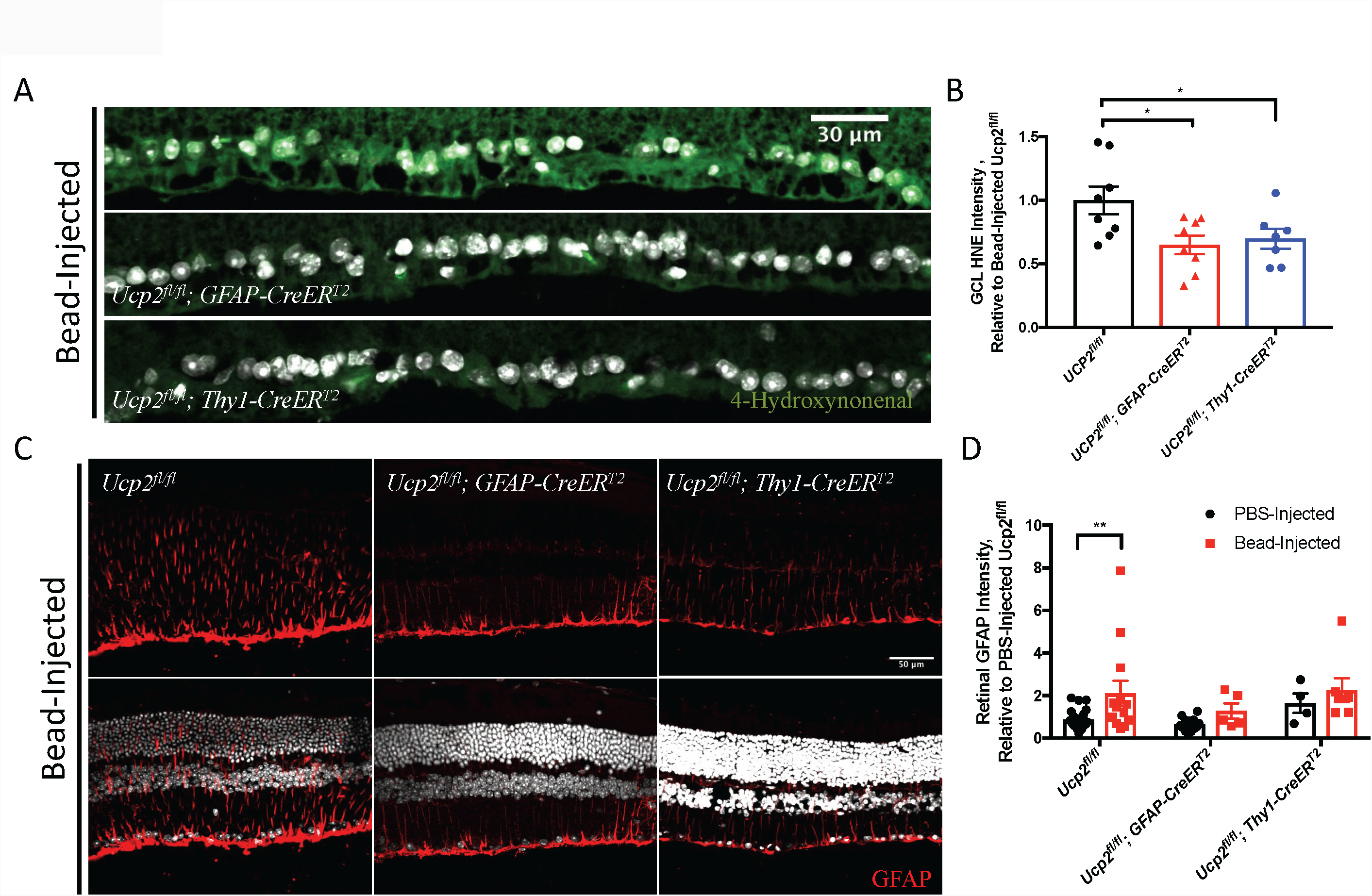
*Ucp2* deletion in glia or retinal ganglion cells decreases retinal oxidative stress and suppresses gliosis. (**A**) Representative 4-hydroxynonenal (green) and Hoechst33258 (blue)-labeled images and (**B**) quantification of 4-hydroxynonenal labeling in the inner retinas of bead-injected *Ucp2*^*fl/fl*^ (n=8), *Ucp2*^*fl/fl*^; *GFAP*-*creER*^*T2*^ (n=7), and *Ucp2*^*fl/fl*^; *Thy1-creER*^*T2*^ eyes (n=8). These data show that the increase in 4-hydroxynonenal observed in glaucoma is decreased by Ucp2-deletion in either RGCs or GFAP-expressing astrocytes and müller glia. The scale bar in (A) is 30 µm. (**C**) Representative images of GFAP labeling of müller glia and astrocytes of bead-injected *Ucp2*^*fl/fl*^ (n=23), *Ucp2*^*fl/fl*^; *GFAP*-*creER*^*T2*^ (n=13) and *Ucp2*^*fl/fl*^; *Thy1-creER*^*T2*^ (n=7) retinas. (**D**) Quantification of retinal GFAP labeling intensity corresponding to (C). GFAP intensity is significantly increased following bead-injection in *Ucp2*^*fl/fl*^ mice, but not in *Ucp2*^*fl/fl*^; *GFAP*-*creER*^*T2*^ or *Ucp2*^*fl/fl*^; *Thy1-creER*^*T2*^ *mice.* Scale bar = 50 µm. *=p<0.05,**=p<0.01.

Neuroprotection of RGCs in transgenic mice was associated with a decrease in HNE labeling intensity, and we hypothesized that Ucp2 deletion altered mitochondrial dynamics or physiology in such a way as to reduce mitochondrial oxidant production. We labeled untreated (n=5) and 4-hydroxytamoxifen (1 µM)-pretreated (n=4) primary astrocytes from *Ucp2*^*fl/fl*^; *GFAP-creER*^*T2*^ mice for the translocase of the outer mitochondrial membrane (Tomm20) to visualize mitochondrial dynamics (Fig. 6A). We quantified mean mitochondrial size, number, mass, and network size using the ImageJ plugin MiNA V1 (Valente et al., 2017). There was a non-significant trend towards decrease in overall mitochondrial mass per cell (p=0.31, df=7, Fig. 6B), a significant decrease in the number of mitochondria/cell (p=0.035, df=7, Fig. 6C), and an increase in mean area (p=0.0061, df=7, Fig. 6D). The overall number of mitochondrial networks/cell was unaltered. Together the images of these mitochondria together with the quantified data suggest and a slight increase the number of more fused mitochondria (Fig. 6A). A healthy pool of mitochondria is maintained through a balance of mitochondrial autophagy (mitophagy) and biogenesis (Piantadosi et al., 2008; Uittenbogaard and Chiaramello, 2014; Ito and Di Polo, 2017). The decrease in mitochondrial number could result from a decline in mitochondrial biogenesis, or an increase in the disposal of damaged mitochondria. We tested the first of these hypotheses by determining the expression of three genes involved in mitochondrial biogenesis, *PolgA, Tfam,* and *PGC1-α.* These genes were not significantly altered by *Ucp2* deletion in primary *Ucp2*^*fl/fl*^; *GFAP-creER*^*T2*^ astrocytes, though the expression of nuclear and mitochondrial genes associated with mitochondrial function (*CytB, Sod2)* were significantly decreased (p=0.037 and 0.024 respectively, df=27, n=3-4, Fig. 6E).

**Figure 6.**
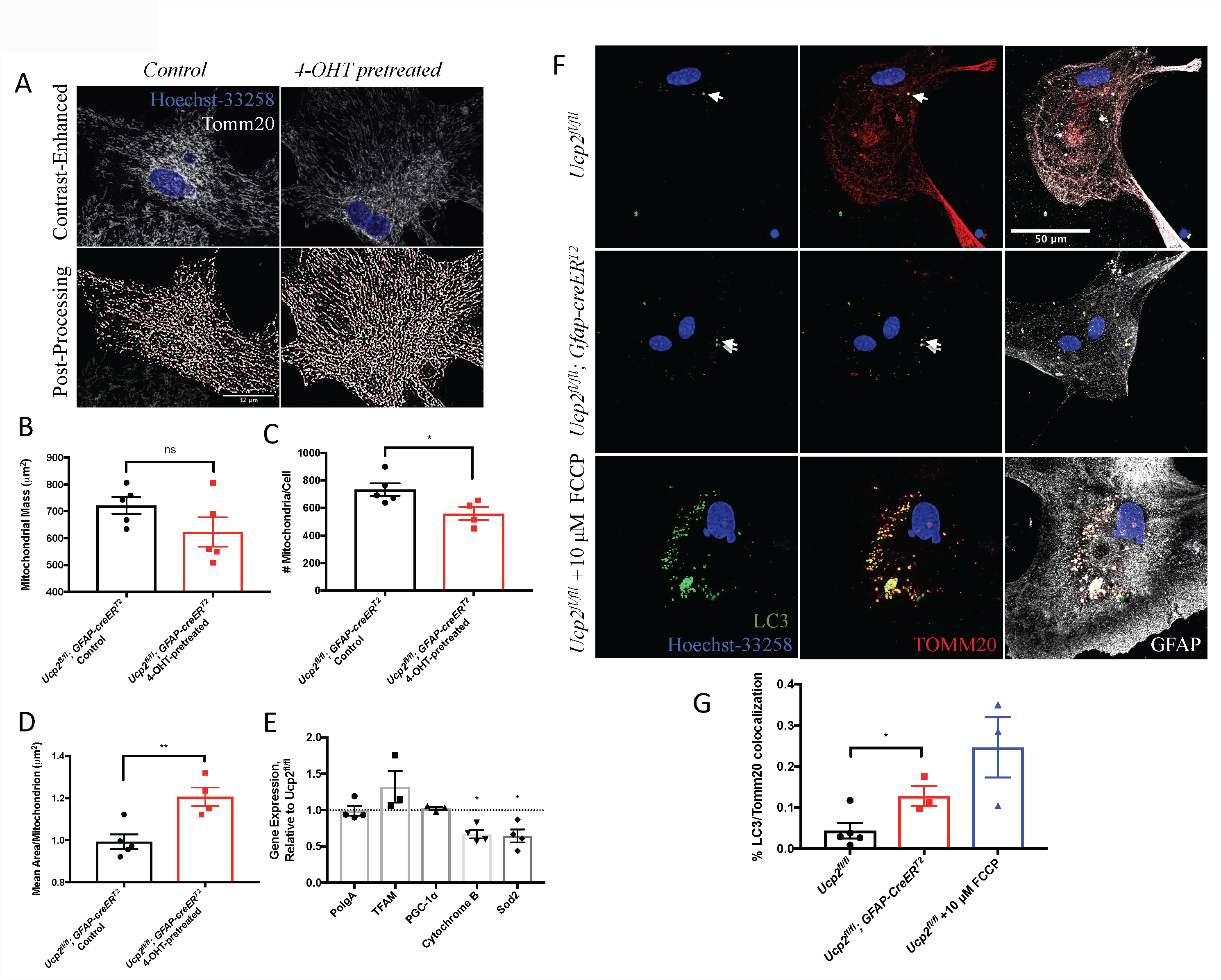
*Ucp2* deletion does not alter mitochondrial biogenesis and increases mitophagy. (**A**) Contrast-enhanced and auto-segmented images of Hoechst-33258 (blue)- and Tomm20 (white)-labeled mitochondria in control and 4-hydroxytamoxifen pretreated *Ucp2*^*fl/fl*^; *GFAP*-*creER*^*T2*^ primary cortical astrocytes (**B-E**) Quantification of morphological morphology through measurement of (**B**) Overall area per cell (mitochondrial mass, µm^2^), (**C**) the number of mitochondria per cell, and (**D**) mean area per mitochondrion (µm^2^) (n=5 biological replicates, derived from different mice with ≥6 cells per biological replicate). Scale bar = 32 µm. (**E**) Expression of genes associated with mitochondrial biogenesis and function in cultured astrocytes, showing that *Ucp2* deletion does not increase biogenesis, but does decrease the expression *Sod2* and *cytochrome B* (n=4). Representative images (**F**) and quantification of colocalization (**G**) between Lc3b (green) and Tomm20 (red) in *Ucp2*^*fl/fl*^ (n=5) and *Ucp2*^*fl/fl*^; *Gfap-creER*^*T2*^ astrocytes (n=3), showing a significant increase in colocalization with *Ucp2* deletion. Cell identity was confirmed using the astrocyte marker GFAP (white) and the DNA label Hoechst-3328 (blue). Image scale is the same as in (A).

We next tested whether the decrease in mitochondrial number and increase in size resulted from activation of mitophagy. We measured the subcellular distribution of the autophagosome adapter protein Lc3b and the mitochondrial outer membrane marker Tomm20 in primary cultures of cortical astrocytes (Fig. 6F-G). Using the Coloc2 plugin on ImageJ, we determined the Manders’ overlap coefficient corresponding to the proportion of co-localized (Lc3b+ Tomm20+) pixels over Tomm20+ pixels (tM2). *Ucp2* deletion significantly increased tM2 from 0.044±0.019 to 0.128±0.024 (p=0.033, df=6, n=3-4, Fig. 6G). tM2 also increased to a greater extent in *Ucp2*^*fl/fl*^ astrocytes treated with 10 µM FCCP for 3h, a treatment known to reproducibly induce mitochondrial fragmentation and mitophagy (0.246±0.074).

To recapitulate these findings in tissue, we reevaluated mitochondrial biogenesis in retinal tissue from *Ucp2*^*fl/fl*^, *Ucp2*^*fl/fl*^; *GFAP-creER*^*T2*^, and *Ucp2*^*fl/fl*^; *Thy1-creER*^*T2*^ mice, by measuring the expression of *PolgA, Tfam,* and *PGC1-α.* Similar to our findings in primary cortical astrocytes, we found no differences in the expression of these factors (Fig. 7A). We also measured the expression of several factors related to mitophagy in the retinas of bead- and PBS-injected transgenic mice (Fig. 7B, n=3/group). We observed genotype- and condition-specific increases in the expression of *Pink1, Park2, Bnip3L*, and *Lc3b*. Specifically, there was an overall significant increase in *Pink1* expression in *Ucp2*^*fl/fl*^; *Thy1-creER*^*T2*^ mice regardless of bead injection (p=0.038), a general increase in *Park2* expression in *Ucp2*^*fl/fl*^; *GFAP-creER*^*T2*^ mice, which was only significant following bead injection (p=0.016). *Bnip3l* was non-significantly elevated in *Ucp2*^*fl/fl*^; *GFAP-creER*^*T2*^ retinas, but only with PBS-injection (p=0.087). Conversely, *Bnip3l expression* was significantly elevated in the retinas from Bead-injected *Ucp2*^*fl/fl*^; *Thy1-creER*^*T2*^ mice but not in PBS-injected controls (p=0.039). *Lc3b* expression significantly increased in bead-injected *Ucp2*^*fl/fl*^; *GFAP-creER*^*T2*^ retinas (p=0.02) and reached a near significant increase in *Ucp2*^*fl/fl*^; *Thy1-creER*^*T2*^ retinas (p=0.058). The expression results suggest that there is an increase in retinal mitophagy following *Ucp2*-deletion, but the specific cell type and environment from which *Ucp2* is deleted will dictate which components of mitophagy machinery are altered.

**Figure 7.**
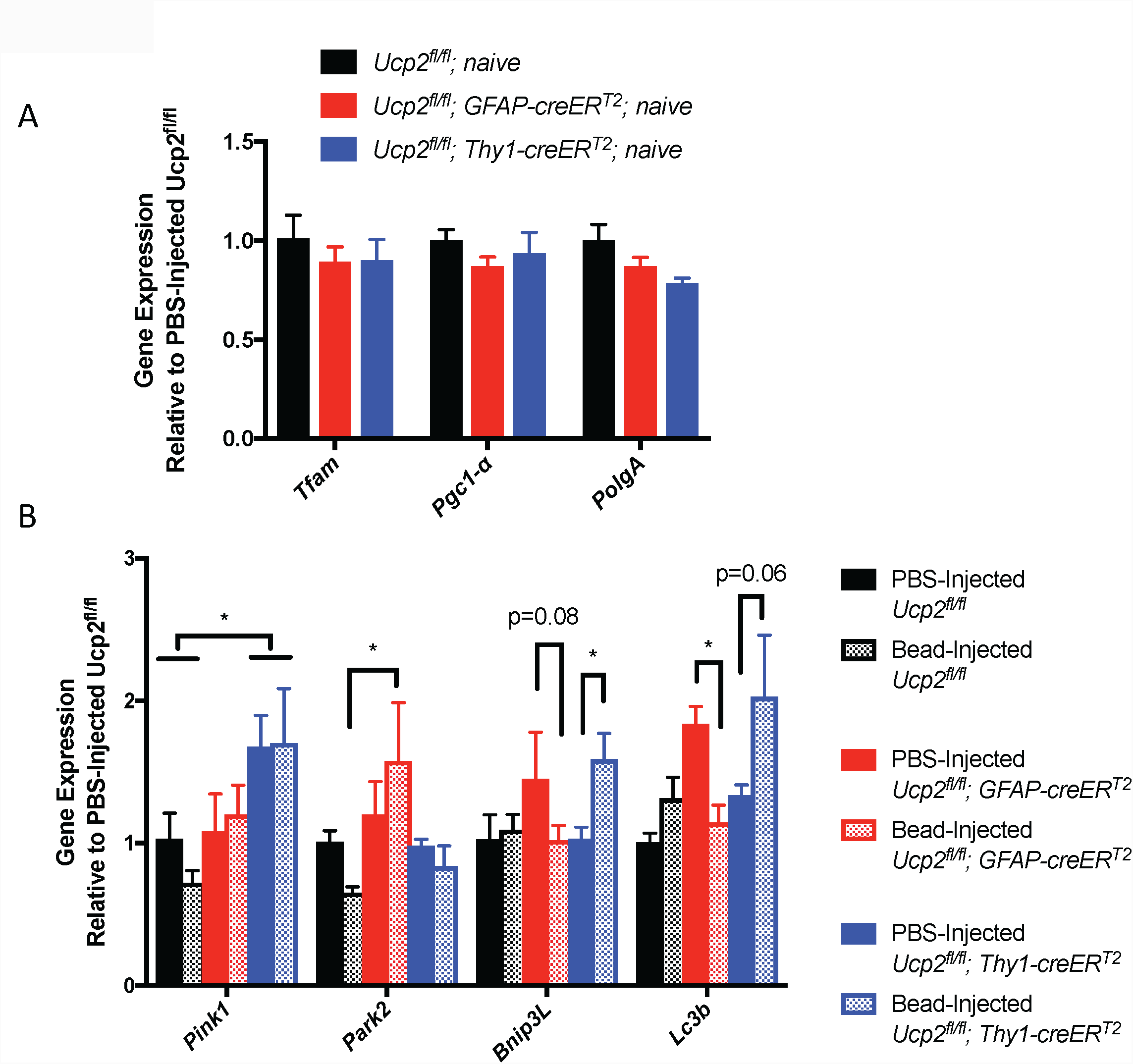
Ucp2 deletion increases the expression of mitophagy-related genes without altering mitochondrial biogenesis. (**A**) Expression of the mitochondrial biogenesis-related genes in retinas from *Ucp2*^*fl/fl*^, *Ucp2*^*fl/fl*^; GFAP-*creER*^*T2*^, and *Ucp2*^*fl/fl*^; *Thy1-creER*^*T2*^ mice. There was no significant change in expression of these genes between mouse lines, though there appears to be a trend towards a decrease in the expression of these factors in the retinas of *Ucp2*^*fl/fl*^; GFAP-*creER*^*T2*^, and *Ucp2*^*fl/fl*^; *Thy1-creER*^*T2*^ mice (n=3). (**B**) Expression of the mitophagy related genes *Pink1, Park2, Bnip3l*, and *Lc3b* in *Ucp2*^*fl/fl*^; GFAP-*creER*^*T2*^ and *Ucp2*^*fl/fl*^; *Thy1-creER*^*T2*^ retinas from PBS-injected (black) and bead-injected (red) eyes, relative to *Ucp2*^*fl/fl*^ control retinas (n=3). 2-way ANOVAs were performed for each gene, accounting for transgene type of transgene and type of injection (PBS or Bead) as independent variables. Bead injection alone did not have a significant effect on any of these genes, genetic background significant impacted *Pink1* expression, and the interaction between bead-injection and genetic background for *Lc3b* and *Bnip3l* expression. A post hoc analysis of each gene revealed other differences in expression, labeled in the figure. *=p<0.05, **=p<0.01, ***p<0.005.

We measured protein levels of both nonlipidated (Lc3b-I) and lipidated (Lc3b-II) forms of Lc3b (Fig. 8A), and found a significant increase in Lc3b-II within both PBS and bead-injected retinas *Ucp2*^*fl/fl*^; *Thy1-creER*^*T2*^ retinas (p=0.029 and 0.0015, respectively, df=15, n=3-4/group, Fig. 8B). We also noticed an increase in total Lc3b levels in the PBS- and bead-injected retinas of both *Ucp2*^*fl/fl*^; *GFAP-creER*^*T2*^ (p=0.0002 and 0.014 respectively, df=15, n=3-4) and *Ucp2*^*fl/fl*^; *Thy1-creER*^*T2*^ (p=0.021 and 0.005 respectively, df=15, n=3-4) mice (Fig. 8C). When we studied the effect of Ucp2-deletion on Bnip3l (Fig. 8D), we found a significant increase in Bnip3l within both transgenic strains (p=0.033, df=14, n=3-4 for *Ucp2*^*fl/fl*^; *GFAP-creER*^*T2*^ and p=0.033, df=14, n=3-4 for *Ucp2*^*fl/fl*^; *Thy1-creER*^*T2*^), but this increase was exclusive to PBS-injected control eyes (Fig. 8E). Notably, the most clearly defined bands in Bnip3l blots migrated at 76 kDa and 100 kDa, respectively corresponding to the Bnip3l dimer (Chen et al., 2010) and a nonspecific band. Analysis of protein levels was performed exclusively on the 76 kDa band, though analysis of the whole lane below 100 kDa yielded almost identical relative protein density values.

**Figure 8.**
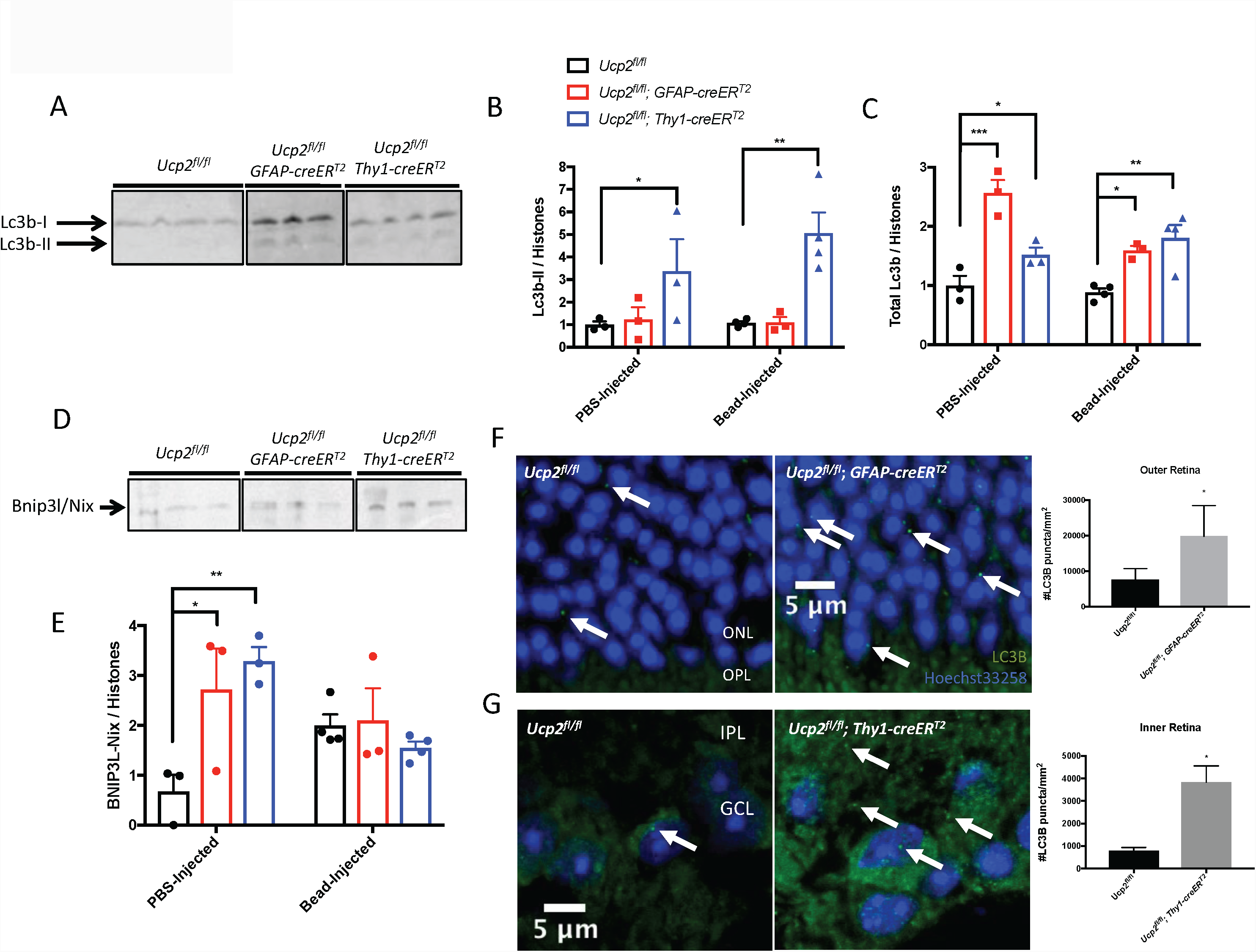
Ucp2 deletion increases Lc3b protein and retinal mitophagy. (**A,D**) Western blot of protein extracts from PBS-injected *Ucp2*^*fl/fl*^ (black bars, n=3-4/group), *Ucp2*^*fl/fl*^; *GFAP*-*creER*^*T2*^ (red bars, n=3/group) *and Ucp2*^*fl/fl*^; *Thy1*-*creER*^*T2*^ (blue bars, n=3-4/group) retinas, labeled for Lc3b-I and Lc3b-II (**A**) as well as Bnip3l/Nix (**D).** Corresponding quantification of protein levels for both PBS- and Bead-injected retinas are shown in (**B**-**C**,**E**). **(B)** Lc3b-II was significantly elevated in both PBS- and bead-injected *Ucp2*^*fl/fl*^; *Thy1*-*creER*^*T2*^ retinas. (**C**) Total Lc3b levels were significantly elevated in PBS- and bead-injected retinas from both *Ucp2*^*fl/fl*^; *Thy1*-*creER*^*T2*^ and *Ucp2*^*fl/fl*^; *GFAP*-*creER*^*T2*^ mice. (**E**) Bnip3l/Nix was significantly elevated in PBS injected retinas of both cre-expressing transgenic mice, but not with bead injection. (**F,G**) We also labeled Lc3b (green) and DNA (Hoechst-33258, blue) in retinal tissue from *Ucp2*^*fl/fl*^ (n=4), *Ucp2*^*fl/fl*^; *GFAP*-*creER*^*T2*^ (n=3), and *Ucp2*^*fl/fl*^; *Thy1-creER*^*T2*^ (n=3) mice. Example images are on the right, and quantifications of Lc3b puncta density are on the left. Compared to the same regions of *Ucp2*^*fl/fl*^ retinas, Lc3b puncta significantly increase in the outer retinas of *Ucp2*^*fl/fl*^; *GFAP*-*creER*^*T2*^, and in the inner retinas of *Ucp2*^*fl/fl*^; *Thy1*-*creER*^*T2*^ mice. *=p<0.05, **=p<0.01, ****=p<0.001.

The RNA and protein expression data yield differing accounts of the molecular events that occur following neural or glial *Ucp2* deletion, and we wanted to determine whether these expression changes led to an increase in autophagosomes in retinal tissue. Within this tissue, mitochondria are too dense for an accurate colocalization analysis of a mitochondrial marker with Lc3b, so we determined the density of autopagosomes in retinas from bead-injected *Ucp2*^*fl/fl*^, *Ucp2*^*fl/fl*^; *GFAP-creER*^*T2*^, and *Ucp2*^*fl/fl*^; *Thy1-creER*^*T2*^ mice. Retinal tissue from Ucp2-deleted mice generally labeled more strongly than *Ucp2*^*fl/fl*^ tissue, which is consistent with an increase in total Lc3b protein. In addition to the increase in background Lc3b levels, bead-injected transgenic retinal tissue had significantly more Lc3b puncta relative to bead-injected Ucp2^fl/fl^ controls. The regional distribution of the increase in these puncta was unique for each cre variant. The density of Lc3b+ autophagosomes increased to a greater extent within the outer retina in *Ucp2*^*fl/fl*^; *GFAP-creER*^*T2*^ mice (p=0.044, df=5, n=3-4, Fig. 8F), though there were non-significant increases in the inner retinas of these mice which may be more clearly resolved with a larger number of replicates. Conversely in bead-injected *Ucp2*^*fl/fl*^; *Thy1-creER*^*T2*^ retinas, the increase in Lc3b puncta density was restricted to the inner retina, which contains RGC soma, dendrite, and axon mitochondria (p=0.01, df=4, n=3, Fig. 8G). Together, the data suggest that Ucp2 deletion increases mitochondrial autophagy in the retina.

## Discussion

Mitochondrial dysfunction and oxidative stress are central to the pathophysiology of glaucoma and have been observed in both animal models and human tissue (Moreno et al., 2004; Malone and Hernandez, 2007; Lee et al., 2011; Mousa et al., 2015; Khawaja et al., 2016). Following a microbead injection, we confirmed the role of elevated IOP in RGC death (Fig. 1), mitochondrial dysfunction, and oxidative stress (Fig. 2).

Previous studies have shown that greater levels of *Ucp2* expression are protective against various conditions that increase oxidative stress, including excitotoxicity and MPTP-induced parkinsonism (Diano et al., 2003; Andrews et al., 2005; Barnstable et al., 2016). We have found that UCP2 is highly expressed in the inner retina (Fig. 1A), an area greatly affected by oxidative stress in glaucoma (Izzotti et al., 2003; Moreno et al., 2004), and have verified the role of Ucp2 in regulating cellular Ψ_m_ and oxidative stress (Fig. 3C-D). Müller glia, astrocytes, and RGCs all exist within the inner retina and each regulate the bioenergetic and oxidative status of ganglion cells (Kawasaki et al., 2000), which is why we modeled glaucoma in mice where Ucp2 was selectively depleted in *Gfap*-expressing (astrocytes and müller glia) or in *Thy1*-expressing cells (retinal ganglion cells; Fig. 4A-B).

We increased the IOP of these mice by injecting microbeads and found that while an elevated IOP may increase RGC loss, reactive gliosis and oxidative retinal damage in *Ucp2*^*fl/fl*^ control mice, *Ucp2*^*fl/fl*^; *GFAP*-*creER*^*T2*^ and *Ucp2*^*fl/fl*^; *Thy1-creER*^*T2*^ were protected from these phenotypes (Fig. 4–5). This finding, while not unprecedented (de Bilbao et al., 2004), was surprising, as *Ucp2* overexpression is neuroprotective in retinas exposed to excitotoxic factors (Barnstable et al., 2016) and the reverse genetic manipulation was expected to have the opposite effect. To determine the source of the observed neuroprotective phenotype, we monitored the mitochondrial morphology of Ucp2-deficient cells. The mitochondria of these cells appeared to be in a more fused state (Fig. 6A-D) and exhibited an elevated respiratory capacity (Fig. 3E), suggesting that mitochondrial function was bolstered by Ucp2 deletion. This led us to investigate the role that *Ucp2* plays in regulating cellular and retinal mitochondrial dynamics.

Health of the mitochondrial population is maintained in part by mitochondrial biogenesis, which can generate new and undamaged mitochondria, and also by mitochondrial autophagy (also known as mitophagy (Lemasters, 2005)), which selectively disposes of damaged or dysfunctional mitochondria, which are smaller and have generally undergone fission (Kim et al., 2007; Frank et al., 2012). Our data did not suggest that Ucp2 altered the transcription of factors associated with mitochondrial biogenesis, either in cells (Fig. 6E) or in retinal tissue (Fig. 7A). We then measured mitophagy in astrocytes, through the association of mitochondrial and autophagosomal proteins Tomm20 and Lc3b. We found a positive indication that mitochondrial components were being degraded more in Ucp2-deficient cells (6F-G), which was well supported by a previous study of *Ucp2* deletion in pulmonary endothelial cells (Haslip et al., 2015). Our cellular data was generally supported by increases in the expression of several mitophagy-related genes *in vivo* (Fig. 7B), though notably these expression profiles did not always correspond with increases in mitophagy or autophagy protein markers. Transcriptional profiling is infrequently used to monitor mitophagy, but the discrepancy between gene expression and gene product levels are still important for studies aiming to translate their findings to the clinic by altering the regulation of mitophagy. In that regard, we do show that *Ucp2* levels may regulates expression of mitophagy genes.

We supported the gene expression data by measuring protein levels of Lc3b-II, a lipidated form of the protein which is anchored in autophagosome membranes (Tanida et al., 2004), as well as the mitochondrial membrane-integral Lc3b adaptor Bnip3l (also known as Nix), a more selective mark of mitophagy rather than generalized autophagy (Sandoval et al., 2008; Chen et al., 2010; Novak et al., 2010; Johansen and Lamark, 2011). Ucp2 deletion in *Thy1*-expressing neurons had a particularly large impact on protein levels of Lc3b-II, and it is unclear why the same effect was undetectable in *Ucp2*^*fl/fl*^; *GFAP*-*creER*^*T2*^ retinal tissue, particularly given that levels of Bnip3l do increase in this tissue (Fig. 8D-E), and the density of Lc3b+ autophagosomes also increases in immunolabeled retinal sections (Fig. 8F-G). These data however are based on tissues with exposed to elevated IOPs for different periods of time. Protein expression data is based on tissue isolated 3 days post-bead injection and Lc3b puncta counts were from tissue isolated 30 days post-bead injection, and together the data suggest that while there autophagic and mitophagic markers are always elevated in *Ucp2*^*fl/fl*^; *Thy1-creER*^*T2*^ retinas, their expression is likely much more dynamic in *Ucp2*^*fl/fl*^; *GFAP*-*creER*^*T2*^ retinal tissue. Of note is the finding that total Lc3b levels are always elevated, and if the stimulation of mitophagy in müller glia or astrocytes is dynamic, that dynamism is likely supported by a larger pool of available Lc3b protein.

The observed increase in mitophagy stimulated by Ucp2-deletion is likely a source of neuroprotection. Mitophagy is already decreased in the glaucomatous mouse optic nerve (Coughlin et al., 2015), and Ucp2-deletion may be protective by re-balancing baseline levels of mitophagy. Similarly, overexpression of the E3 ubiquitin ligase Parkin in the rat retina stimulates retinal mitophagy and is also protective against glaucomatous retinal ganglion cell death (Dai et al., 2018). Therefore, the RGC-intrinsic increases in mitophagy following Ucp2 deletion are probably directly protective, though our investigations in *Ucp2*^*fl/fl*^; *GFAP*-*creER*^*T2*^ mice lead us to the critical question of how glial mitophagy can protect RGCs from oxidative stress and cell death.

Retinal glia are phagocytic (Bejarano-Escobar et al., 2017), and astrocytes of the optic nerve head are able to dispose of damaged RGC mitochondria under physiological conditions (Davis et al., 2014), which leads us to the hypothesis that a Ucp2-deletion dependent increase in glial mitophagy increases the transcellular disposal of RGC mitochondria. To determine whether this is the case in future studies; we will need a sophisticated methodology that combines transgenic Ucp2 deletion in glia with a fluorescent probe targeted to retinal ganglion cell mitochondria. With this setup we could determine whether modulation of glial *Ucp2* increases the density of RGC mitochondria engulfed by glia.

Our data point to a non-canonical function for *Ucp2,* and perhaps for uncoupling proteins in general. While they decrease oxidative damage when active, uncoupling protein inactivity may promote a sub-lethal form of oxidative damage that triggers mitophagy (Frank et al., 2012), specifically selecting dysfunctional mitochondria for degradation (Lemasters, 2005; Kim and Lemasters, 2011). This activity could result in cells enriched for undamaged mitochondria, which we see when imaging mitochondria. This model is represented in Figure 9. Overall, we suggest that as a consequence of regulating mitochondrial ROS homeostasis, *Ucp2* is also a regulator of mitochondrial dynamics.

**Figure 9.**
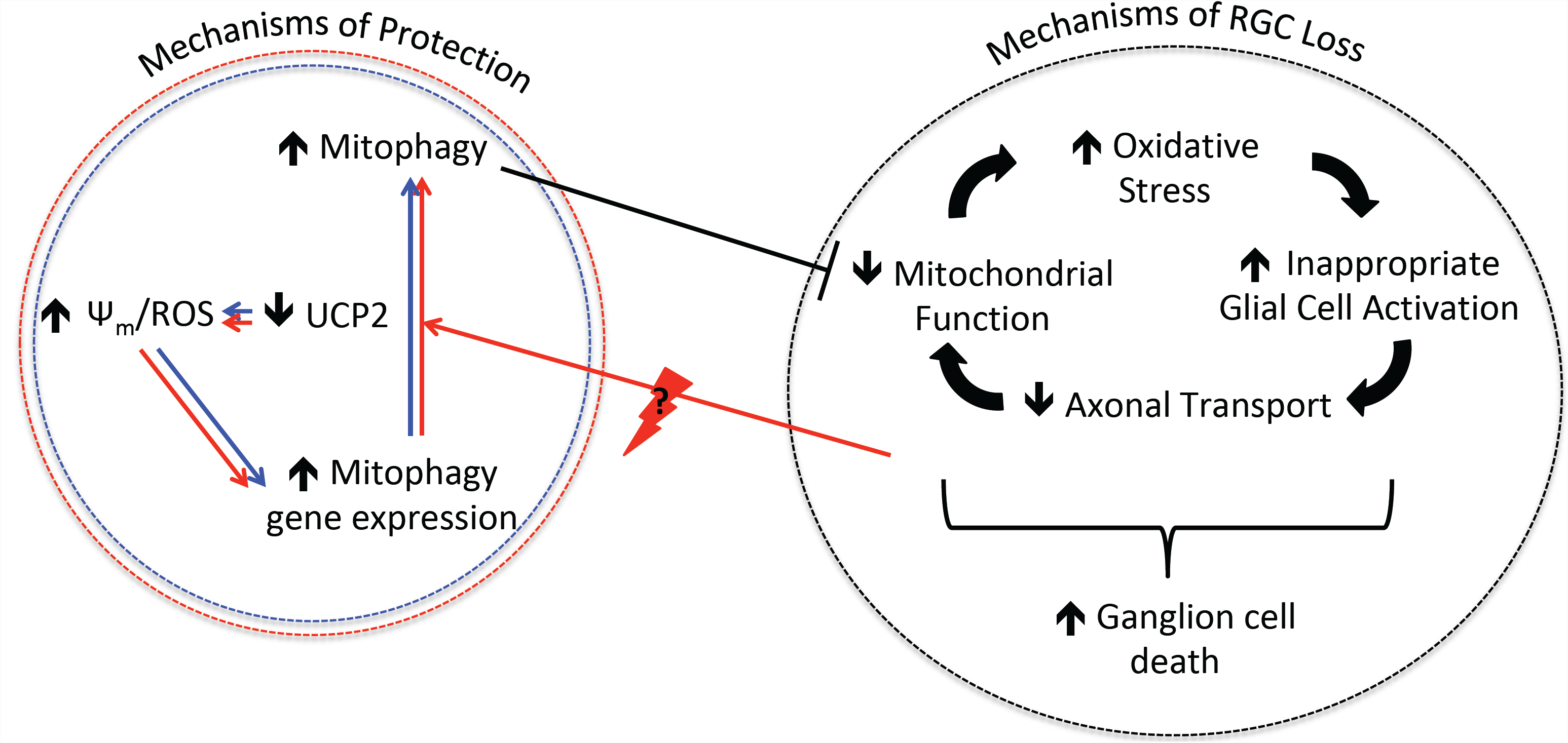
The proposed model for Ucp2 deletion-mediated neuroprotection. Schematic diagram showing the interplay of oxidative stress, decreased mitochondrial function, decreased axonal transport, and altered glial cell activation, all of which are known to play pathophysiological roles in glaucoma and several other neurodegenerative conditions. These factors all contribute to each other and to retinal ganglion cell death. Our proposed mechanism to explain the protective effects of *Ucp2* deletion is by this alteration leading to a Ψ_m_ and ROS-dependent increase in mitophagy, occurring both through increases in mitophagy gene expression and changes in protein levels, which provide a pool of mitophagy components and increase the dispoal of damaged mitochondria either at baseline when *Ucp2* is deleted in RGCs (blue) or in inducibly in *Gfap*-expressing glia (red). The stimulus which causes the more dynamic changes in glial mitophagy is represented by a thunderbolt enclosing a “?”. In this model, the increase in mitophagy cuts short degenerative phenomena that result from dysfunctional mitochondria, and decreases cellular damage.

## Author Contributions Statement

DTH and CJB contributed conception and design of the study; DTH acquired, analyzed, interpreted the data, and wrote the first draft of the manuscript; DTH and CJB contributed to critical revision and final approval of the manuscript.

## Acknowledgments

This work was supported by grants from the National Institutes of Health and the Macula Vision Research Foundation. The authors thank Angela Snyder for providing the mouse cartoon used in figures 1 and 3, as well as Drs Evgenya Popova, Gregory Yochum, and Ian Simpson for their critical review of and intellectual contributions to this manuscript.

